# Transposable Element Evolution in the Allotetraploid *Capsella bursa-pastoris*

**DOI:** 10.1101/042325

**Authors:** J. Arvid Ågren, Hui-Run Huang, Stephen I. Wright

## Abstract

**Premise of the study:** Shifts in ploidy affect the evolutionary dynamics of genomes in a myriad of ways. Population genetic theory predicts that transposable element (TE) proliferation may follow because the genome wide efficacy of selection should be reduced and the increase in gene copies may mask the deleterious effects of TE insertions. Moreover, in allopolyploids TEs may further accumulate because of hybrid breakdown of TE silencing. However, to date the evidence of TE proliferation following an increase in ploidy is mixed, and the relative importance of relaxed selection vs. silencing breakdown remains unclear.

**Methods:** We used high-coverage whole genome sequence data to evaluate the abundance, genomic distribution, and population frequencies of TEs in the self-fertilizing recent allotetraploid *Capsella bursa-pastoris* (Brassicaceae). We then compared the *C. bursa-pastoris* TE profile with that of its two parental diploid species, outcrossing *C. grandiflora* and self-fertilizing *C. orientalis*.

**Key results:** We found no evidence that *C. bursa-pastoris* has experienced a large genome wide proliferation of TEs relative to its parental species. However, when centromeric regions are excluded, we find evidence of significantly higher abundance of retrotransposons in *C. bursa-pastoris* along the gene-rich chromosome arms, compared to C.*grandiflora* and *C. orientalis*.

**Conclusions:** The lack of a genome-wide effect of allopolyploidy on TE abundance, combined with the increases TE abundance in gene-rich regions suggest that relaxed selection rather than hybrid breakdown of host silencing explains the TE accumulation in *C. bursa-pastoris*

## Introduction

A central goal of population and comparative genomics research is to understand what factors drive the evolution of genome size and structure (Gregory, 2005; Lynch, 2007; Alfoldi and Lindblad-Toh, 2013; Koenig and Weigel, 2015). Coupled with decades of information on genome size from flow cytometry across diverse lineages, the growing wealth of whole genome sequence data has revealed how extensive and rapidly genome size and structure can evolve, even among close relatives (Ungerer et al., 2006; Hawkins et al., 2009; Wright and Ågren, 2011; Tenaillon et al., 2011; Leitch and Leitch, 2013; Ågren and Wright 2015; Ågren et al., 2015). Yet our ability to explain this variation remains in its infancy.

Whole genome duplication via polyploidization has long been considered to be a major contributor to genome evolution (Adams and Wendel, 2005; Soltis and Soltis,2012; Hollister 2015; Soltis et al. 2015). Most obviously, polyploidization will cause a direct increase in the total amount of DNA per cell. This initial doubling of genome size has often been followed by a process of diploidization, leading to a pattern of DNA loss over time (Leitch and Bennet, 2004; Lysak et al., 2009; Renny-Byfield et al., 2013; Vu et al., 2015). Furthermore, although the direct role of recent polyploidy on genome size evolution can be investigated and controlled for, most plant species have experienced a history of whole genome duplication in their evolutionary history (Vision et al. 2000; Jaillon et al., 2007; Jiao et al., 2011; Vanneste et al., 2014; Li et al. 2015). A lack of complete information on this history can therefore make it difficult to fully investigate the importance of whole genome duplication events on the evolution of genome size and structure.

In addition to the direct effect of ploidy on genome size, transposable element (TE) proliferation may follow whole-genome duplications due to the masking of deleterious insertions and a reduction of the efficacy of selection across the genome caused by genome redundancy (recently reviewed in Parisod and Senerchia, 2012; Tayalé and Parisod, 2013). Furthermore, host-mediated silencing of TEs may be disrupted in allopolyploids (where an increase in ploidy is due to interspecific hybridization; Madlung et al., 2002; 2005; Kraitshtein et al., 2010; Yaakov et al., 2011). This combination of relaxed selection and a breakdown of silencing mechanisms could potentially drive dramatic evolution of genome structure following whole genome duplication. In particular, gene-dense euchromatic regions with very low TE content may experience major accumulation of TEs in genic regions. Such a mechanism may explain, for example, the dramatic transposable element expansion in the maize genome following whole genome duplication (Schnable et al., 2009; Baucom et al., 2009; Diez et al., 2014).

On the other hand, genome downsizing in polyploids may lead to a net loss of transposable elements during the process of diploidization (Parisod et al., 2010). The empirical evidence of changes in TE abundance following an increase in ploidy is equivocal. An increase in TE copy number was reported in *Nicotiana tabacum* (Petit et al., 2007; 2010; but see Renny-Byfield et al., 2011), whereas *Orobanchegracilis* appears to have experienced TE loss (Kraitschtein et al., 2010). Thus, overall, the factors driving proliferation vs. loss of transposable elements in polyploids remain poorly understood, and the relative importance of relaxed selection vs. silencing breakdown in TE accumulation is not clear.

Here, we use high-coverage whole genome sequence data to evaluate the abundance, genomic distribution, and population frequencies of TEs in the self-fertilizing allotetraploid *Capsella bursa-pastoris. Capsella bursa-pastoris*is a recently derived allotetraploid, with population genomic evidence for genome-wide reduction in the strength of selection on point mutations due to both gene redundancy and its selfing mating system (Douglas et al. 2015), making it an interesting model to examine the early fate of transposable elements following allopolyploidization. We look for evidence of TE proliferation beyond what would be expected from additivity of its two parental diploid species and use patterns of genomic distribution of TE insertions to distinguish the relative importance of relaxed selection vs. silencing breakdown. We discuss our results in light of the literature on the association between polyploidization, mating system, and TE abundance across plant species.

## Material and Methods

### Study system

The genus *Capsella* of the Brassicaceae family consists of four species with varying mating system, ploidy, and geographical distribution (Hurka et al. 2012; Figure 1).Selfing is thought to have evolved multiple times from an ancient progenitor of the diploid (2*n* = 2x = 16) *Capsella grandiflora*, a self-incompatible species restricted to Albania and northwestern Greece. Most recently, *Capsella rubella* diverged from *C. grandiflora* within the last 100,000 years (Foxe et al. 2009; Guo et al. 2009; Slotte et al. 2013; Brandvain et al. 2013). *Capsella orientalis*is thought to have evolved selfing prior to *C. rubella*, also from a *C. grandiflora*-like ancestor. *C. orientalis* and *C. grandiflora* have been inferred to have diverged approximately 930,000 years ago, providing the potential for a longer time period of mating system divergence (Douglas et al. 2015). Whereas *C. rubella* has expanded to a larger Mediterranean distribution, *C. orientalis* is now found in an area spanning Eastern Europe to Central Asia (Hurka et al. 2012). The origin of the world-wide distributed *Capsella bursa-pastoris* long remained elusive, but was recently determined to be an allotetraploid (2*n* = 4x = 32) following a hybridization event between *C. grandiflora* and *C. orientalis* within the last 100,000-300,000 years (Douglas et al. 2015). Consistent with this hybrid origin, a principal component analysis of the types of TEs found in *C. bursa-pastoris* puts it as an intermediate between *C. grandiflora* and *C. orientalis* and of all shared insertions found in two of the three species, the majority are between *C. bursa-pastoris* and either *C. grandiflora* or *C. orientalis*, with very few shared between *C. grandiflora* and *C. orientalis* to the exclusion of *C. bursa-pastoris* (Douglas et al. 2015). In this study, we expand the TE analysis of *C. bursa-pastoris*, to test for an accumulation of TEs following allopolyploid origins.

**Figure 1.**
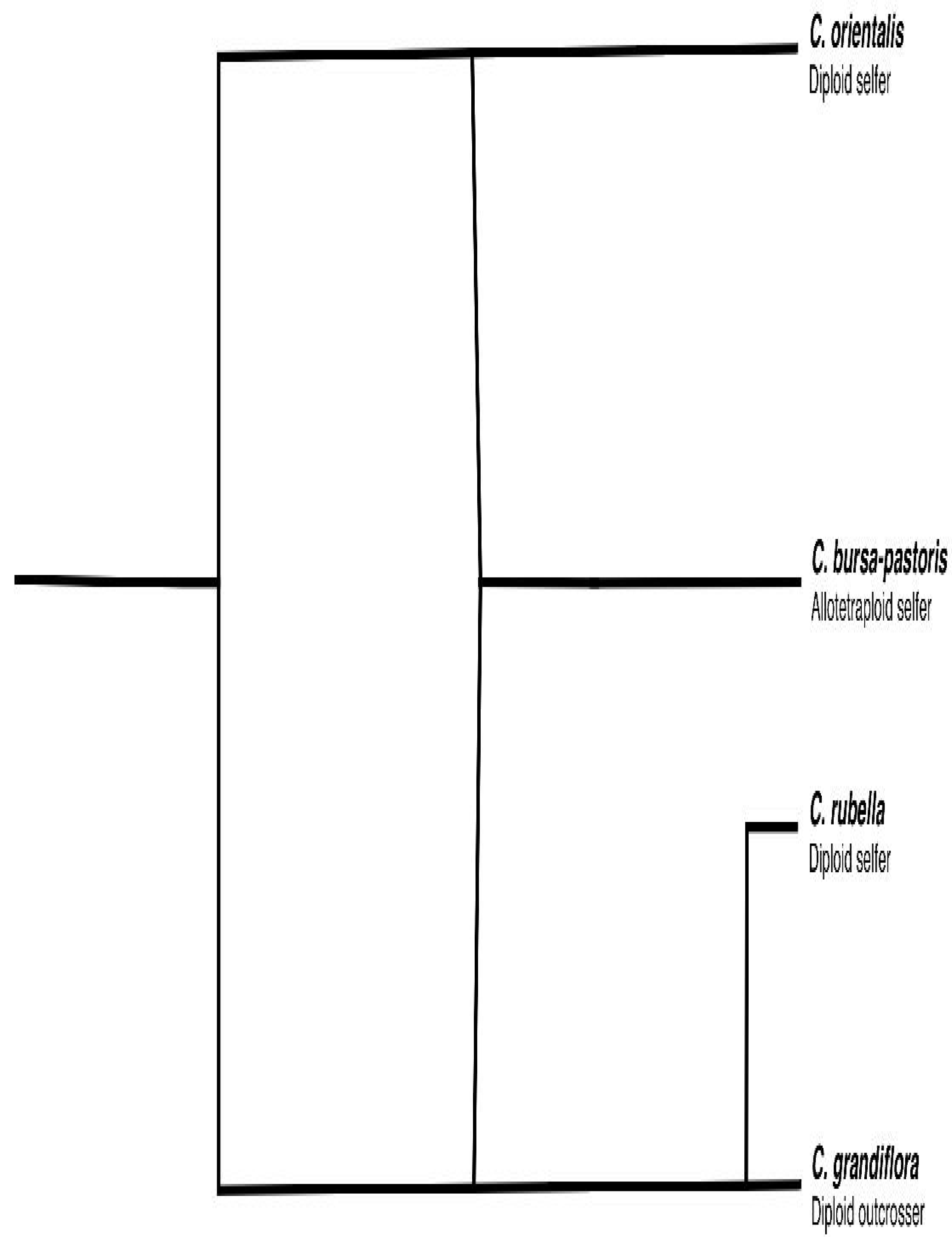
Phylogenetic relationships within the *Capsella* genus. The evolution and natural history of the genus are thoroughly discussed in Hurka et al. (2012), Slotte et al. (2013), and Douglas et al. (2015).

### Identification and quantification of transposable elements

To compare the abundance, genomic locations, and population frequencies of TEs in the three species we combined the TE datasets generated by Ågren et al. (2014) and Douglas et al. (2015). These studies applied the PoPoolationTE pipeline of Kofler et al. (2012) on 108-bp paired-end Illumina reads on 7 *C. bursa-pastoris*, as well as 8 *C. grandiflora* and 10 *C. orientalis* individuals sampled across their respective geographical distributions, which provides the most comprehensive picture of TE evolution in each species. Ågren et al. (2014) also analysed TE distributions in *C. rubella*, but for this study we focus on the two direct progenitor species of *C. bursa-pastoris, C. grandiflora* and *C. orientals.* We used the *C. rubella* reference genome (Slotte et al. 2013) and the TE database generated as part of the same study, which consisted of sequence data from seven Brassicaceae species, including *Capsella rubella, Brassica rapa, Arabis alpina, Arabidopsis thaliana* (accessions Col-0, Ler, Kro-0, Bur-0, and C24 from the 1001 *Arabidopsis* genomes project), *Arabidopsis lyrata, Eutrema halophila*, and *Schrenkiella parvulum* Since the PoPoolation TE approach is designed for pooled population data, the original output is an estimate of the population frequency of each TE insertion. We adjusted the pipeline to use population frequencies to infer insertions as homo-or heterozygous. We ignored as spurious insertions with an estimated frequency of <0.2 and considered insertions with a frequency of >0.8 as homozygous. Intermediate frequency insertions were treated as heterozygous. Note that for our highly selfing tetraploid species, insertions identified as ‘heterozygous' when mapping to the diploid *C. rubella* reference genome are in fact likely to be homozygous in one of the two homeologous genomes, and so for *C. bursa-pastoris*, intermediate frequency insertions were treated as homozygous at one of the two homeologues, while fixed insertions were treated as present in both homeologues. To avoid falsely inferring independent insertions due to the uncertainty in the method in the precise genomic position, we treated insertions as the same if the distance of the inferred location of two or more insertions across individuals was < 200 bp and the inferred TE family was identical. In previous work, we performed extensive tests to ensure that this approach could generally distinguish homo- and heterozygous insertions (Ågren et al. 2014). We used this approach to determine the abundance, genomic locations, and population frequencies of TEs in the three species.

## Results

We quantified the abundance of four major categories of TEs: DNA, Helitrons, long terminal repeat (LTR) retrotransposons and non-LTR retrotransposons. The three species differ in their mean number of TEs (Kruskal-Wallis chi-squared□=□21.342, df□=□2, p□<□0.00001), but genome-wide all species show similar relative abundance across elements, with LTR elements making up the bulk of the insertions (Figure 2).

To test whether *C. bursa-pastoris* has experienced an accumulation or loss of TEs following its origin, we calculated the expected diploid TE copy number from a *C. orientalis × C. grandiflora* hybrid and compared this number to the observed *C. bursa-pastoris* abundance. We randomly paired up *C. orientalis* and *C. grandiflora* chromosomes and calculated the average TE copy number of such a cross (where heterozygous insertions were given a copy number of 0.5). We performed 1,000 replicates of *in silico* crosses, sampling with replacement, based on the present-day TE abundances. We then compared the expected copy number to the observed abundance in *C. bursa-pastoris* and found that *C. bursa-pastoris* harbours slightly but significantly more insertions genome-wide than what would be expected under strict additivity (Figure 2; Wilcoxon rank sum test, p = 0.01297). Thus, overall we do not observe evidence for a major reduction in host silencing driving high rates of transposition.

**Figure 2.**
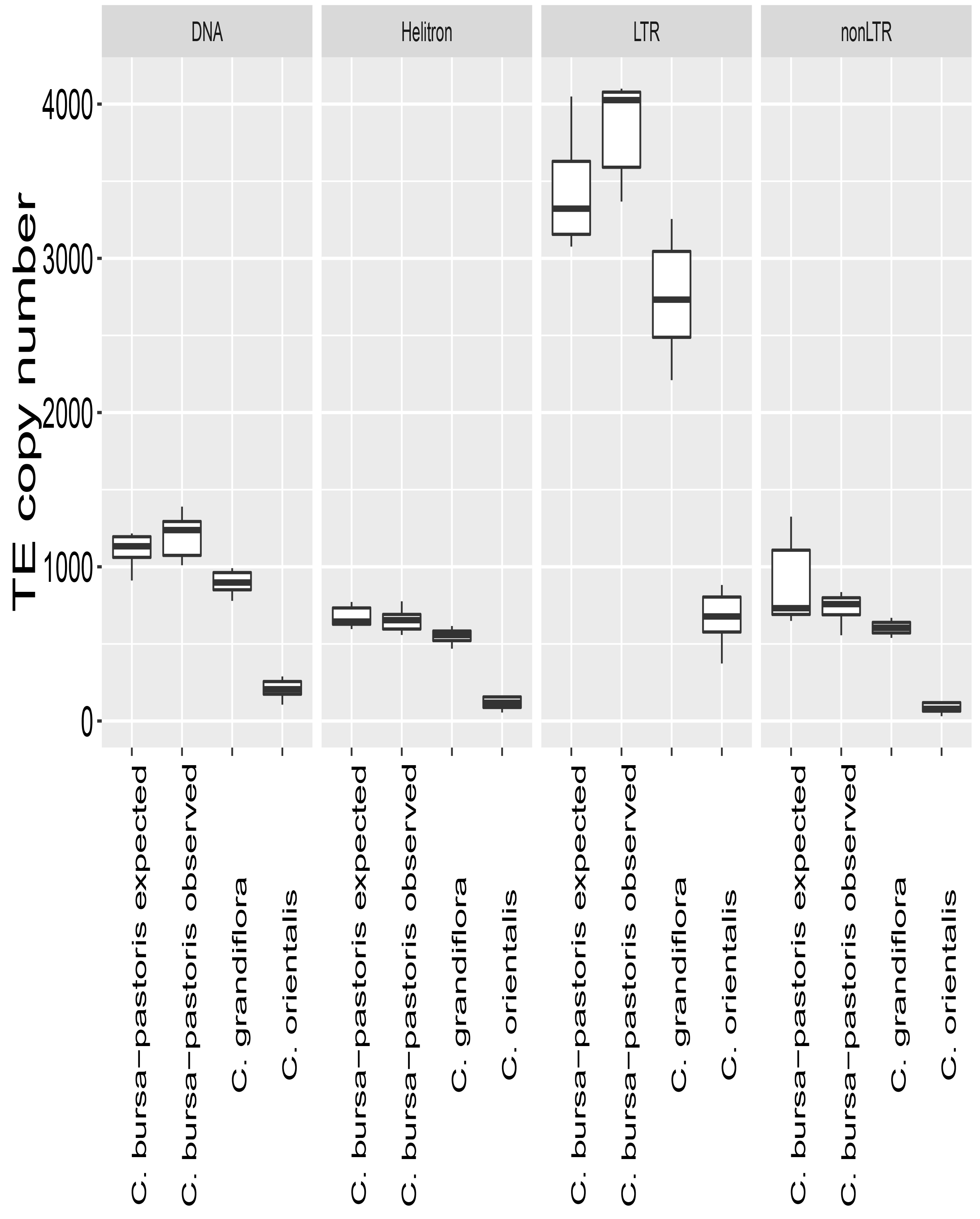
Boxplots of the genome wide abundance of transposable elements (TEs) in the three *Capsella* species. The expected *C. bursa-pastoris* value was generated by performing 1,000 replicates of in silico crosses between *C. orientalis* × *C. grandiflora*, sampling with replacement.

Since TE insertions near genes will likely disrupt gene function, population genetic theory predicts that selection will rapidly remove such insertions (Dolgin and Charlesworth 2008). Following a whole-genome doubling event, a tetraploid like *C. bursa-pastoris* will carry twice as many gene copies as its diploid progenitors and the fitness cost of an insertion should therefore be less. As a consequence, tetraploids may be expected to accumulate more TEs near genes than diploids. To test this prediction, we first excluded the centromeric regions of the genome, following the annotation of Slotte et al. (2013), and considered TE abundance in the gene-rich chromosome arms only. Restricting our attention to these regions, we find that *C. bursa-pastoris* has a considerably higher TE abundance than expected from additivity (Figure 3; Wilcoxon rank sum test, p = 0.0005828), particularly for retrotransposons. Second, we used the gene annotation from the reference genome of *C. rubella* (Slotte et al. 2013) to calculate the distance to the closest gene for all TE insertions, in all three species. Again, just like the overall abundance, we are interested in whether *C. bursa-pastoris* has more insertions near genes than what would be expected by additivity from a *C. orientalis × C. grandiflora* cross. Using the approach outlined above, we calculated the expected TE copy number within 1000 bp of the closest gene from such a hybrid and compared it to the observed abundance in *C. bursa-pastoris.* We find that *C. bursa-pastoris* harbours significantly more insertions near genes, compared to what would be expected under strict additivity (Figure 4; Wilcoxon rank sum test, p = 0.000126).

**Figure 3.**
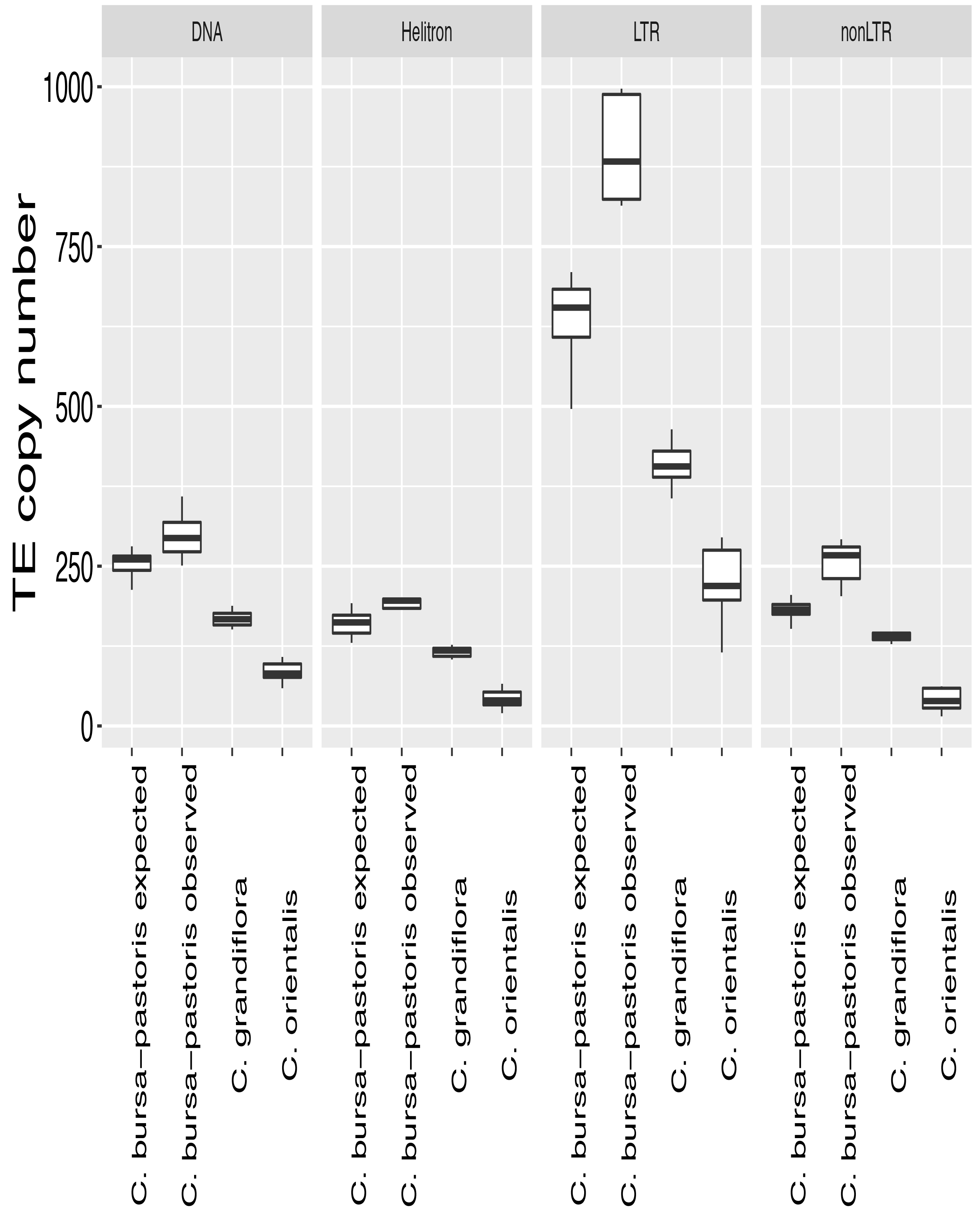
Boxplots of the abundance of transposable elements (TEs) in the three *Capsella* species in the gene rich chromosome arms (centromeric regions excluded). The expected *C. bursa-pastoris* value was generated by performing 1,000 replicates of in silico crosses between *C. orientalis* × *C. grandiflora*, sampling with replacement.

**Figure 4.**
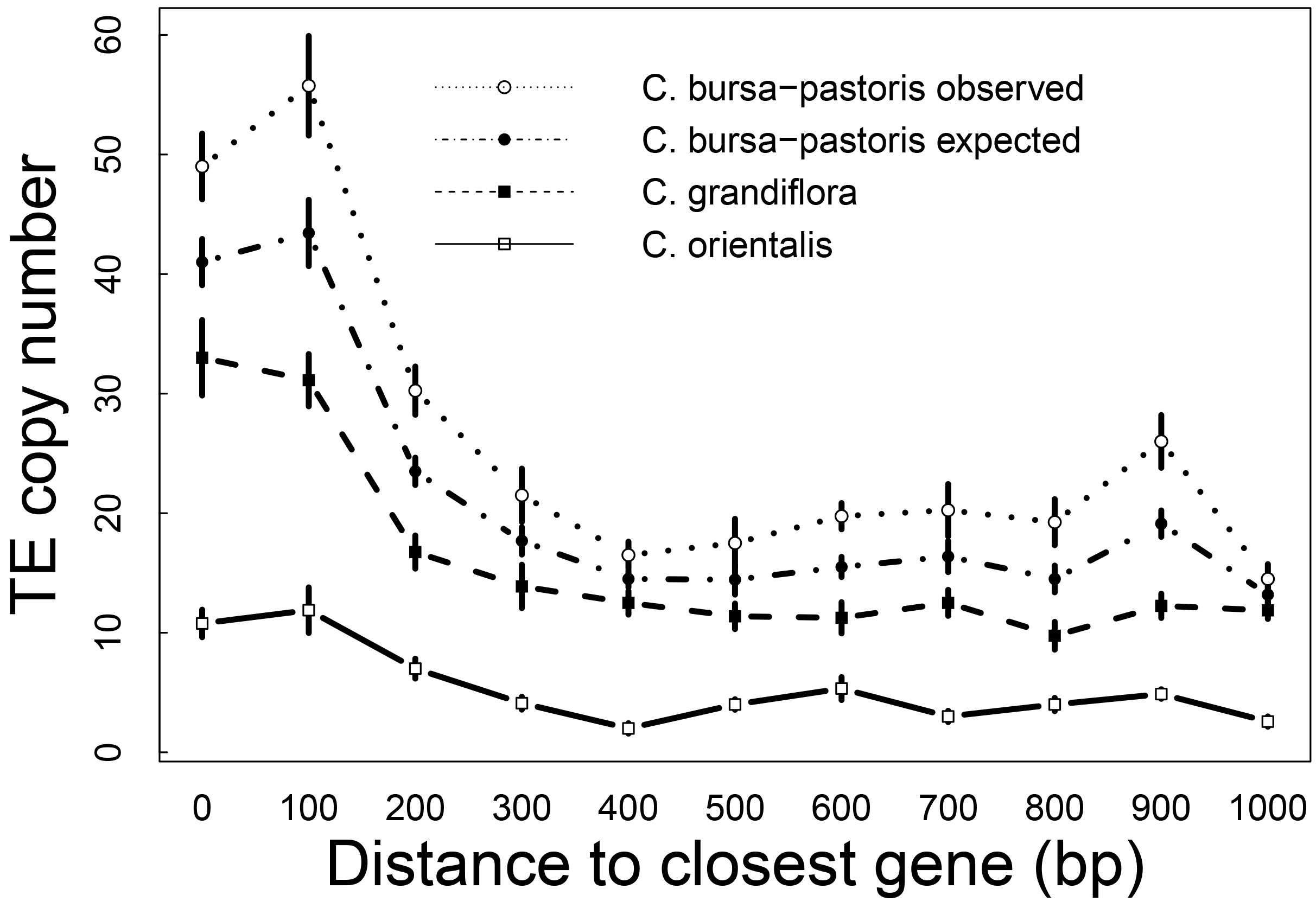
Average abundance of transposable elements (TEs) in 100 bp bins near their closest gene in the three *Capsella* species. Error bars are□±□1 standard error. The expected *C. bursa-pastoris* value was generated by performing 1,000 replicates of in silico crosses between *C. orientalis* × *C. grandiflora*, sampling with replacement.

We used the presence/absence of all TE insertions, across all individuals in the three species to categorize insertions as either singletons (present in only one individual) or non-singletons (present in more than one individual). We find that *C. grandiflora* has the highest proportion of singletons, potentially suggesting a higher TE activity and/or stronger purifying selection against insertions than *C. orientalis* and *C. bursa-pastoris*, which both show similar proportions of singletons (Table 1). However, differences in demographic history between the species are likely also contributing to the frequency spectrum, and the overall count of rare insertions is highest in *C. bursa-pastoris*. Overall, the combination of elevated copy number and a lower proportion of singletons in the tetraploid species is consistent with relaxed purifying selection following the transition to tetraploidy.

**Table 1.** Number of singleton and non-singleton transposon insertions in three *Capsella* species

|  | Singletons | Non-singletons |
| --- | --- | --- |
| *C. orientalis* | 524 (20%) | 2095 (80%) |
| *C. grandiflora* | 3060 (29%) | 7493 (71%) |
| *C. bursa-pastoris* | 3768 (22%) | 13358 (78%) |

To investigate in more detail the relative importance of inherited TE insertions from parental species vs. new ongoing transposition events, we calculated the percentage of *C. bursa-pastoris* insertions found in each diploid progenitor species, separated by insertion frequency class (Figure 5). Note that for insertions found in counts greater than 7 (our sample size), this would imply that the insertion is found in both homeologous genomes at that position. As expected, low-frequency insertions are rarely found in the progenitor diploid species, suggesting a significant number of new insertions in the tetraploid, although clearly some of these cases may have been unsampled in the population but present in an ancestral diploid genome. In contrast, common insertions and those found on both homeologues tend much more often to be found in one or both diploid parental species, reflecting the ‘parental legacy’ (Buggs et al. 2014) of some insertions inherited in the tetraploid. Also as expected, more insertions are generally shared with *C. grandiflora*, reflecting their greater abundance, however a number of intermediate frequency insertions likely reflect fixed insertions found on the C. orientalis homeologue (Figure 5, insertion frequencies of ‘7’ reflect fixed insertions on one homeologue). Even intermediate-frequency insertions however have a large fraction unsampled from diploid progenitors, suggesting relaxed selection on de novo insertions is allowing TEs to spread since polyploid origins. Overall, these patterns highlight both the additive, inherited contribution and the role of new de novo insertions in gene-rich regions in the TE complement of a recent tetraploid.

**Figure 5.**
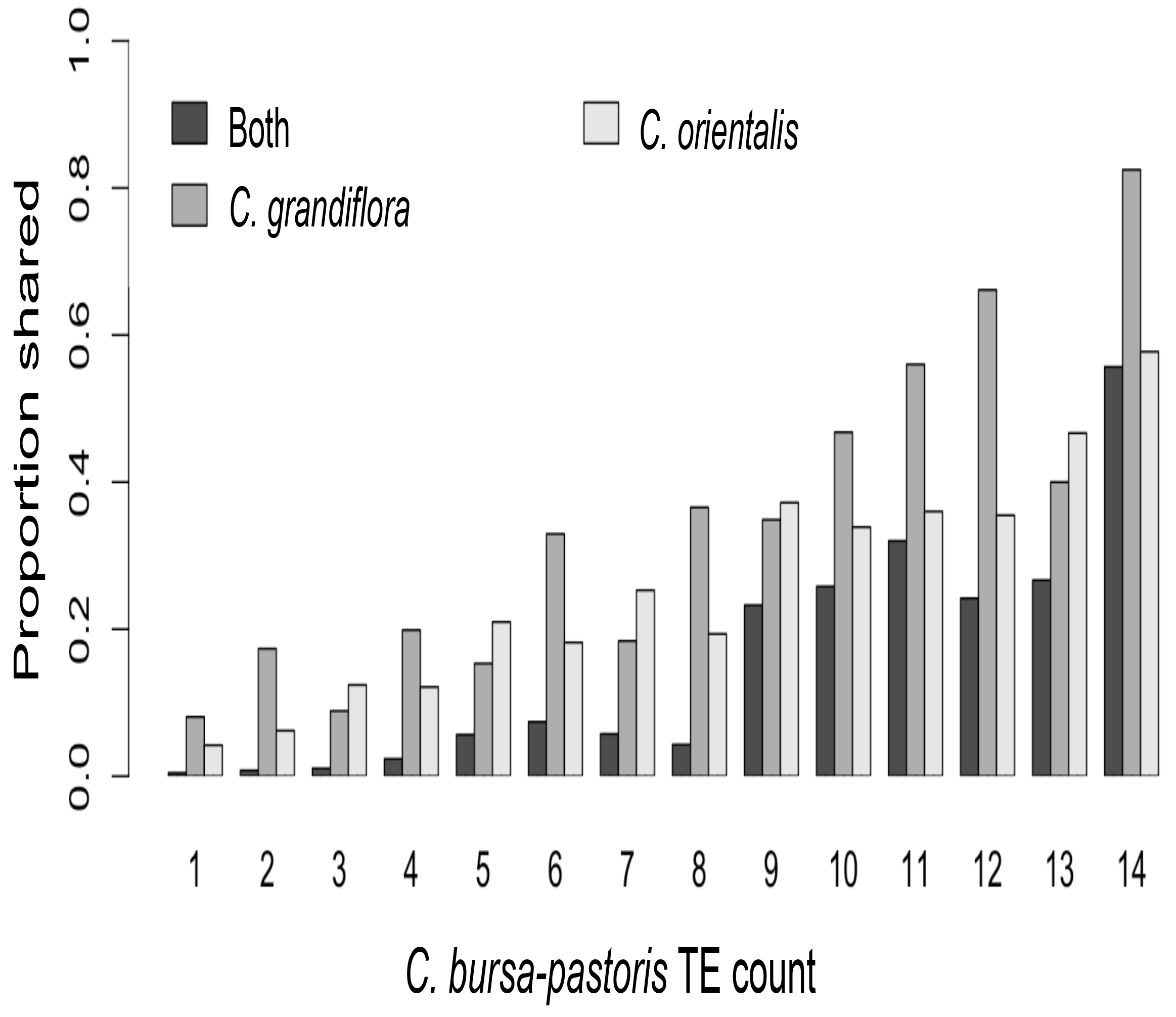
Proportion of *C. bursa-pastoris* transposable element (TE) insertions found in the diploid progenitors *C. orientalis, C. grandiflora*, or both. Proportions are separated by TE frequency class in *C. bursa-pastoris*, where the counts represent the number of copies of the insertion in a sample size of 7 *C. bursa-pastoris* individuals. Note that counts greater than 7 imply the insertion is found on both homeologous copies of the genome, and counts of 7 are likely to be cases of fixation events in one of the two homeologues.

## Discussion

Overall, we found no evidence that *C. bursa-pastoris* is experiencing a large-scale genome wide proliferation of TEs, as would be expected if there were a genome-wide breakdown of host silencing mechanisms. This is consistent with genome size estimates of the species and its prognitors, which does not indicate a non-additive increase in genome size (Hurka et al., 2012). However, we did detect a considerably higher abundance of TEs than expected when restricting our analysis to gene rich regions. These results are in line with previous work by Douglas et al. (2015) on genome-wide SNP patterns, which suggested that while this allopolyploid has not experienced a large-scale ‘genome shock’ since its origin, it is undergoing a global quantitative reduction in the efficacy of selection on amino acid and conserved noncoding mutations. Taken together, our results suggest that long-term relaxation of selective constraints is leading to TE accumulation in gene-rich regions, without a major shift in transposition rate.

One important consideration when predicting the effects of polyploidization on genome evolution may be its association with mating system. Under a number of population genetic models highly outcrossing species are predicted to experience higher rates of transposable element activity and copy number (Wright and Schoen, 1999; Morgan, 2001; Charlesworth and Wright, 2001). Allopolyploidization events that are associated with a retention of high rates of outcrossing could therefore represent a ‘perfect storm’, whereby TE activity remains high while genome redundancy enables rapid proliferation. On the other hand, polyploidy is often associated with elevated rates of self-fertilization compared with diploid relatives, and both asexual reproduction and high rates of selfing are common in polyploid lineages (reviewed in e.g. Mable, 2004; Husband et al., 2008; Robertson et al., 2011; Ramsey and Ramsey, 2014). Highly selfing lineages such as *C. bursa-pastoris* may thus experience a more modest increase in copy number than outcrossers; it would be of interest to investigate similarly-aged outcrossing allopolyploid lineages to assess whether TE accumulation is more dramatic in these species. On the other hand, selfing tetraploid lineages are more likely to experience severe founder events during polyploid origins, and strong genetic drift may further contribute to relaxed selection following whole-genome duplication as we observed in *C. bursa-pastoris* (Douglas et al., 2015).

It is notable that some of the most well-documented ancient TE expansion events, including maize (Schnable et al., 2009; Baucom et al., 2009; Diez et al., 2014) and the *Brassica* genus (Zhang and Wessler, 2004), are associated with ancient allopolyploidization events involving outcrossing lineages. Whether this is simply circumstantial or causal will require in-depth comparative analyses of the joint and unique effects of polyploidy and mating system on genome size and TE proliferation. Although the current age distribution of retroelements in the maize genomes suggests that TE proliferation was more recent than whole genome duplication (Bennett and Leitch 2005), this does not rule out ongoing TE accumulation due to relaxed selection over millions of years. As we gain increasingly detailed insights into the time since last whole genome duplication event in many lineages, investigating how this interacts with mating system to structure genome evolution is becoming increasingly feasible.

## Acknowledgements

We thank two anonymous reviewers for their helpful comments on the manuscript. J.A.A was supported by a Junior Fellowship from Massey College, S.I.W. by a Natural Sciences and Engineering Research Council (NSERC) Discovery Grant, and H.R.H by the National Natural Science Foundation of China (grant number 31370005) and a fellowship from the China Scholarship Council.

